# Maturation-informed synthetic Magnetic Resonance Images of the Developing Human Fetal Brain

**DOI:** 10.1101/2024.04.08.588566

**Authors:** Hélène Lajous, Andrés le Boeuf Fló, Pedro M. Gordaliza, Oscar Esteban, Ferran Marqués, Vincent Dunet, Mériam Koob, Meritxell Bach Cuadra

## Abstract

Magnetic resonance imaging is a powerful modality to investigate abnormal developmental patterns *in utero*. However, since it is not the first-line diagnostic tool in this sensitive population, data remain scarce and heterogeneous between scanners and centers. In addressing the data scarcity issue while generating data representative of real fetal brain MRI, we proposed *FaBiAN*, a ‘Fetal Brain magnetic resonance Acquisition Numerical phantom. Here, we present a novel synthetic dataset of 594 two-dimensional, low-resolution series of T_2_-weighted images corresponding to 78 developing human fetal brains between 20.0 and 34.8 weeks of gestational age. Data are generated with substantive improvements from the original *FaBiAN* to account for local heterogeneities within white matter tissues throughout maturation. These synthetic-yet-highly-realistic images cover both healthy and pathological development trajectories simulated with standard clinical settings and anatomically informed by the Fetal Tissue Annotations (FeTA) dataset. Two independent radiologists qualitatively assessed the realism of the simulated images. We also quantitatively demonstrate the simulated data’s increased fidelity to real data compared to the previous *FaBiAN* version. The reuse potential of the proposed dataset was also evaluated in the context of automated fetal brain tissue segmentation. Besides, our dataset that combines images generated from various clinical scenarios has been made publicly available to support the continuous endeavor of the community to develop advanced post-processing methods aswell as cutting-edge artificial intelligence models.

## Background & Summary

There is a growing awareness of the importance of early brain maturation on health later in life as the underlying complex, interconnected structural and functional processes can be altered by various genetic and environmental factors^1–9^. Accurate characterization of *in utero* development is therefore critical.

Magnetic resonance imaging (MRI) is an emergent adjunct to ultrasound (US) in cases of diagnostic ambiguity, and for comprehensive diagnostic, prognostic, and postnatal management planning^10^. MRI is adequate for exploring the developing fetal brain due to its excellent contrast in soft tissue while being minimally invasive. Clinical guidelines recommend the acquisition of T_2_-weighted (T2w) fast spin echo (FSE) sequences to workaround unpredictable fetal motion during the exam. In practice, at least three orthogonal series of two-dimensional (2D) thick slices are acquired to provide information on the whole brain volume with sufficient signal-to-noise ratio (SNR)^11^. Despite all counter-measures and optimizations, fetal MRI remains challenging due to motion artefacts and related signal drops, as well as low SNR in small structures within the maturing brain surrounded by the mother’s womb and the amniotic fluid. Since MRI is second-line and not the reference-standard technique (i.e, US) for the follow-up of the fetus during pregnancy, large-scale clinical datasets are relatively scarce in this cohort of sensitive subjects. Besides, there is no standardized imaging protocol across sites, which has resulted in large variability between MR schemes across scanners, studies^12^, and even more so between vendors. Such discrepancies may result in highly variable MR contrasts and image quality. Indeed, most of the data available today are heavily post-processed and integrated into spatio-temporal MRI atlases of the fetal brain, either healthy^10,13,14^ or pathological^15^. This enables a fine representation of the developing brain throughout gestation. However, such atlases average brain scans across several fetuses at a given gestational age (GA), thus resulting in high-resolution (HR) images far from a realistic clinical set-up, with smoothed inter-individual heterogeneities and features. Recently, the Fetal Tissue Annotations (FeTA) dataset has been proposed as a benchmark for automated multi-tissue fetal brain segmentation^16,17^. However, only super-resolution (SR) reconstructions^18,19^ of the fetal brain volume and their associated semi-automated annotations have been made publicly available, but not the original clinical acquisitions. In fact, to date, no database provides annotated low-resolution (LR) series of the fetal brain.

Thanks to their ability to provide a flexible and controlled environment that facilitates accurate, robust, and reproducible research, computer simulations are widely used for MR developments to mitigate data scarcity and post-processing complexity^20–26^. In this context, we demonstrated that synthetic, yet realistic data can efficiently complement scarce clinical datasets, providing valuable support fot data-demanding deep learning (DL) models for fetal brain MRI tissue segmentation^26–28^, as well as the optimization of advanced reconstruction techniques^26,29–31^. These exploratory studies were based on the first Fetal Brain magnetic resonance Acquisition Numerical phantom (FaBiAN) that simulates as closely as possible the FSE sequences used in clinical routine for fetal brain examination to generate realistic T2w images of the fetal brain throughout maturation from a variety of segmented HR anatomical images of healthy and pathological subjects^26^. Despite a good tissue contrast, the synthetic T2w MR images used in this work were originally derived from a three-class model of the fetal brain (gray matter (GM), white matter (WM), and cerebrospinal fluid (CSF)) that does not allow to capture key maturation processes and metabolic changes occurring in WM tissues across gestation.

In the wake of this first prototype^32^, the proposed data descriptor showcases a full dataset of highly realistic in silico data composed of:

i. 594 synthetic T2w MR images corresponding to 78 developing fetal brains, derived from HR annotations of SR-reconstructed, real clinical data acquired on various MR scanners and following the different clinical protocols in place at Lausanne University Hospital (CHUV) and at University Children’s Hospital Zurich (Kispi);
ii. automatically-generated brain masks and fetal brain annotations of the LR series;
iii. a corresponding SR reconstruction for every subject.

This dataset is based on a numerical model of the developing fetal brain that accounts for the pronounced MR signal changes representative of tissue heterogeneity within WM structures throughout maturation^33^. Our methodological contribution (FaBiAN v2.0^34^) thus identifies local spatial regions of variable water content within a WM mask to simulate more realistic *in utero* MR images.

Figure 1 provides a schematic overview of the study design presented in this work. We specifically validate the data as follows. First, we report a large-scale, independent, qualitative assessment of the realism of the newly simulated images (FaBiAN v2.0) compared to the former implementation (FaBiAN v1.2) by two neuroradiology and pediatric radiology experts. Second, we compare quantitatively the SR volumes reconstructed from real clinical cases and from their corresponding synthetic LR images generated using FaBiAN v1.2 and v2.0. Finally, we demonstrate a potential use-case of the dataset with an *in silico* augmentation of training samples for automated fetal brain multi-tissue segmentation of *in utero* MR acquisitions, through the simulation of a broad variety of sequence parameters using FaBiAN v2.0.

**Figure 1.**
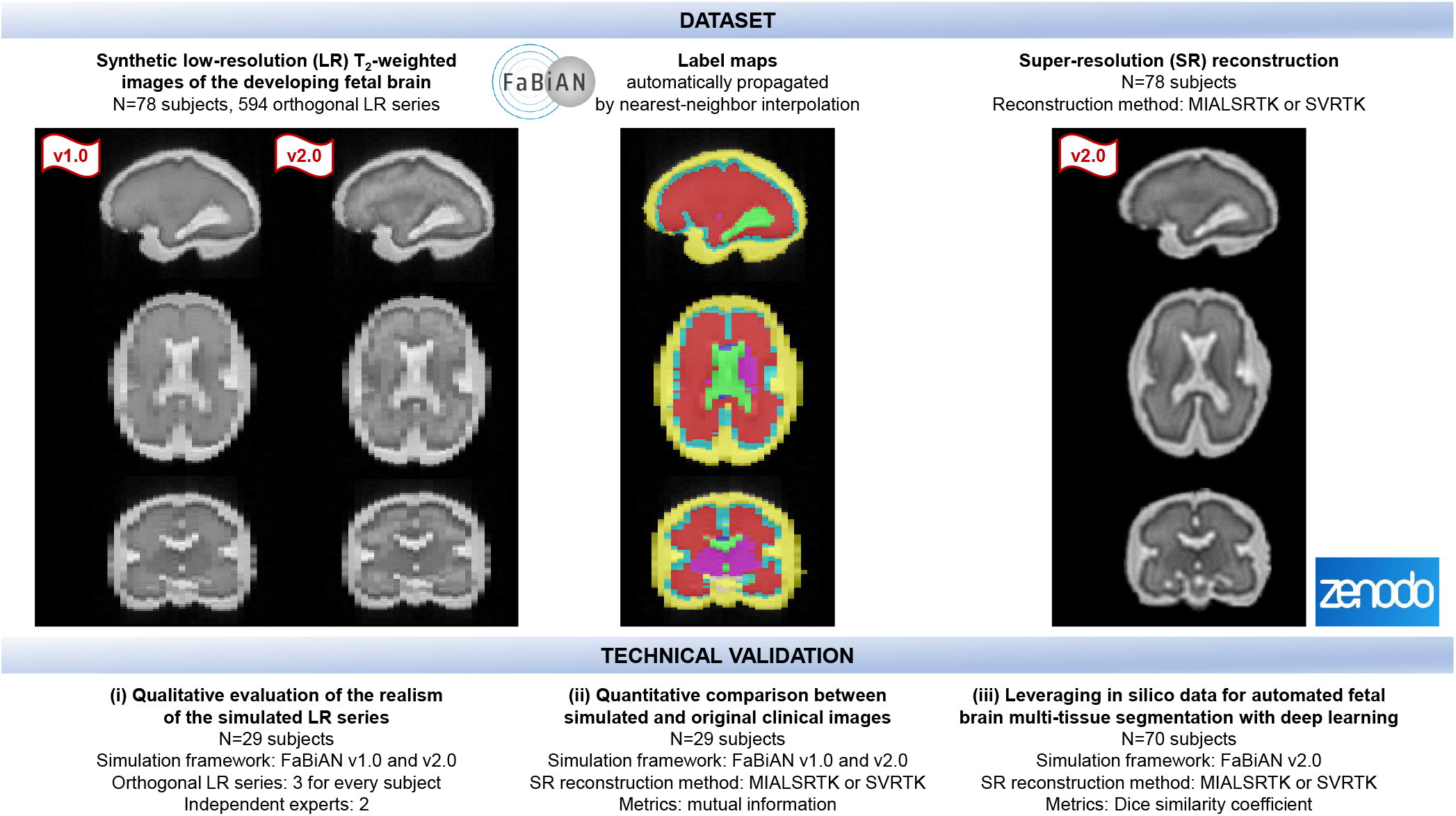
Schematic overview of the dataset promoted throughout this work and of the study design to validate its relevance to the fetal brain MRI community.

This study aims at demonstrating the relevance of such an extensive dataset of synthetic, yet highly realistic T2w MR images of the fetal brain throughout maturation to a community struggling with data scarcity in this sensitive population that requires comprehensive ethical oversight to acquire new data as well as experienced MR technologists. As such, having access to multiple MR images generated from various settings and in different clinical scenarios has a great potential reuse value for further developments of advanced post-processing methods as well as cutting-edge artificial intelligence models.

## Methods

### Data

The input anatomical models used to generate all the LR series of the fetal brain with local WM changes across maturation for the technical validation were extracted from the publicly available FeTA Dataset with refined annotations^35,36^. This clinical dataset gathers 88 subjects (34 neurotypical and 54 pathological, two were excluded after quality control due to the bad quality of the SR reconstruction) in the GA range of 20.0 to 34.8 weeks (27.0 *±* 3.58 weeks). Common pathological conditions such as spina bifida, ventriculomegaly, and heterotopia are included. Eight tissues are segmented: WM, intra-axial CSF, cerebellum, extra-axial CSF, cortical GM, dGM, brainstem, and corpus callosum.

The data used in this study were acquired in earlier studies in accordance with the relevant guidelines and regulations, under the supervision of Ethics Boards composed of representatives at different levels (hospitals, cantons, and federal state). Mothers of all fetuses included in the current work provided written informed consent for the re-use of their data for research purposes.

### Modeling of white matter heterogeneity and changes across maturation

Figure 2 gives an overview of FaBiAN v2.0, our latest developments to account for local WM heterogeneities throughout fetal brain maturation. In particular, Figure 3 highlights the major changes implemented to modulate brain tissue relaxation times across GA.

**Figure 2.**
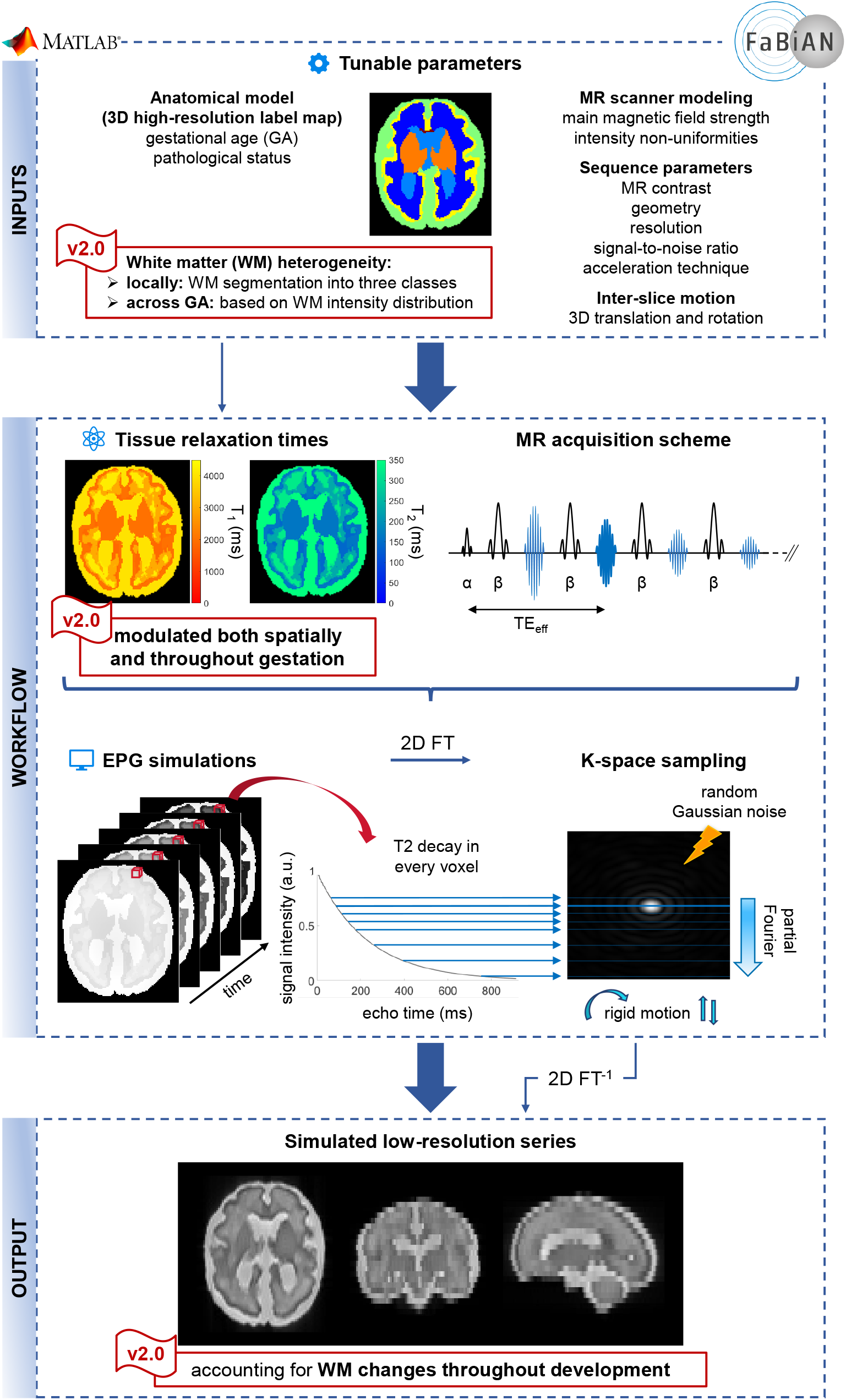
Workflow for simulating fast spin echo (FSE) acquisitions of the fetal brain which better capture local WM changes throughout maturation (FaBiAN v2.0). The major changes compared to the original implementation of the software are highlighted by the flag “v2”.

**Figure 3.**
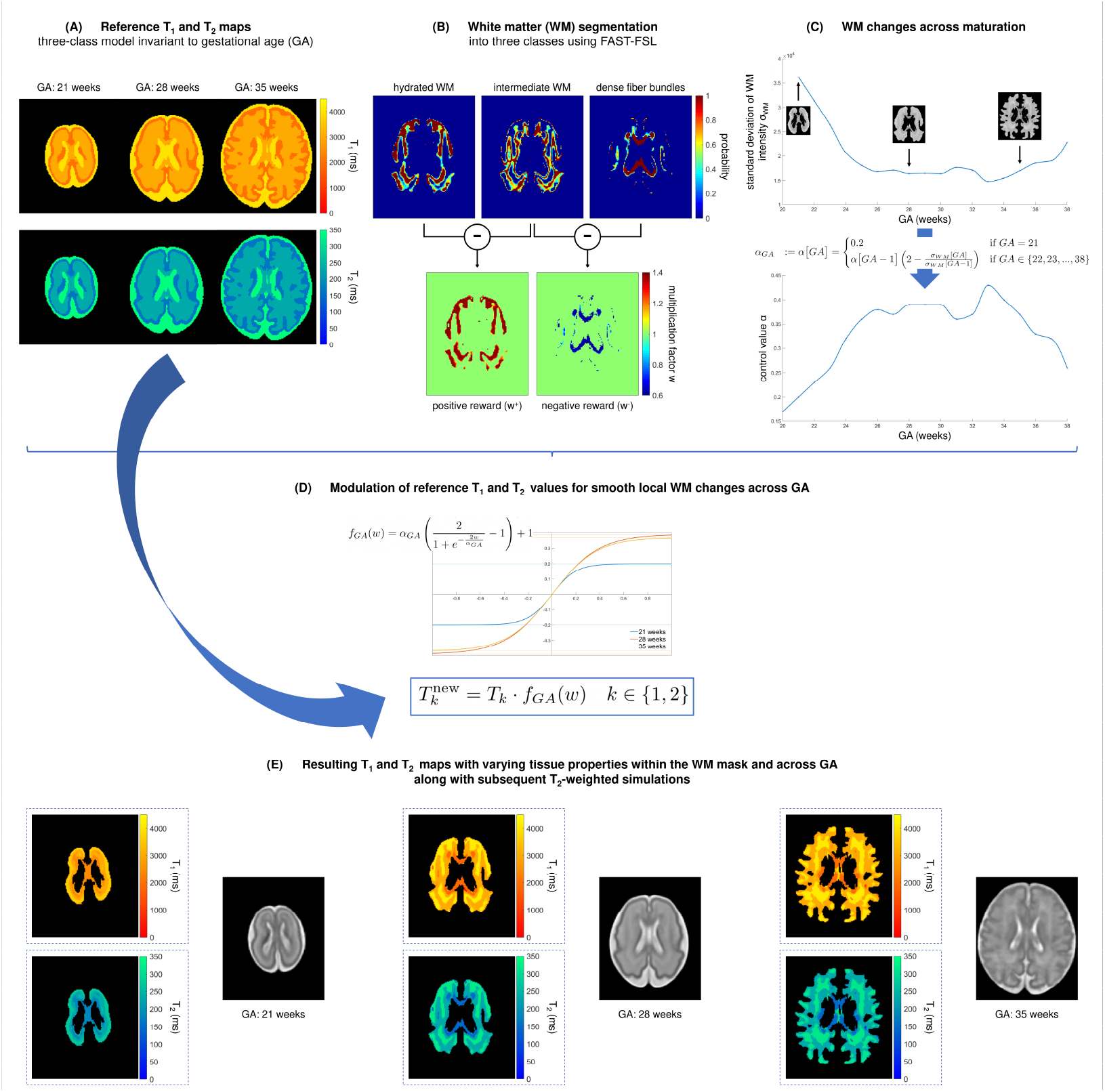
Modulation of reference *T*_1_ and *T*_2_ values in the WM to depict smooth, local WM changes throughout fetal brain maturation.

Due to the lack of characterizations or normative maps of T_1_ and T_2_ relaxometry measurements of fetal brain structures, all tissues from the segmented fetal brain images are merged into three main classes: GM, WM and CSF. According to previous empirical relaxometry values provided in the literature^37–41^, an average T_1_, respectively T_2_ value is assigned to each of these classes, without consideration of the GA or spatial a priori on the finer location within the brain^26^ (see Figure 3-(A)).

We rely on the Gaussian Hidden Markov Random Field (GHMRF) model, fitted using the Expectation-Maximization algorithm, following the approach outlined in FMRIB’s Automated Segmentation Tool (FAST^42^) to integrate local spatial WM heterogeneities in our numerical representation of the developing fetal brain, and therefore capture biophysical changes that arise across maturation. Concretely, we automatically segment a WM mask into three classes using FAST 6.0.5.1 as illustrated in Figure 3-(B)): hydrated WM areas that appear hyperintense on a T2w image, hypointense areas representing dense WM fibers, and a third class interpreted as an intermediate between hydrated WM and maturing, dense WM fibers. Parameters were set heuristically as follows: 0.1 MRF regularization weight; four iterations for bias field removal; 20.0 mm kernel in bias field smoothing.

The segmentation output consists in 3D partial volume (PV) concentration maps for each voxel (*i*) within the three segmented classes. These PV maps are used to ultimately weight T_1_ and T_2_ relaxation times locally. Specifically, the intermediate WM class is considered as a baseline: the positive difference between the PV maps of hydrated, respectively dense WM and this baseline will be used to increase, respectively decrease the average reference T_1_ and T_2_ relaxation times in every voxel of the WM mask. The corresponding weight (*w*^+^, respectively *w*^−^) computed in every voxel of the WM mask according to Equation 1 can be displayed in so-called *positive and negative reward maps* (Figure 3-(B)).

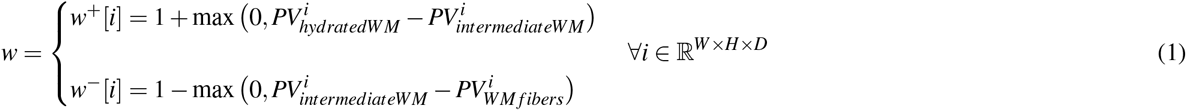

where: *w*^+^ and *w*^−^ are respectively the positive and negative reward maps, *PV*_*hydratedWM*_, *PV*_*intermediateWM*_, and *PV*_*WM fibers*_ stand for the PV concentration maps of each of the three segmented WM classes, respectively the hydrated WM, the intermediate WM class, and the dense WM fiber bundles. *W, H*, and *D* represent the width, height, and depth of the maps respectively.

The adjustment of the mean reference T_1_ and T_2_ relaxation times should be approached with caution though as substantial weights (*w*) may result in unrealistic relaxometric properties of the modeled fetal brain tissues. To address this concern, a sigmoid modulation characterized by a control value *α* (see Figure 3-(D)) is applied to the original T_1_ and T_2_ values in order to smoothen local WM changes within brain structures and ensure that the corresponding tissue properties do not excessively deviate from their reference values. Because the range of WM intensity changes is not uniform across gestation (see Figure 3-(C)), *α* needs to be adjusted according to the GA. We rely on the normative spatiotemporal MRI atlas (STA) of the fetal brain which includes 18 subjects spanning 21 to 38 weeks of GA^10^ to determine the optimal setting following:

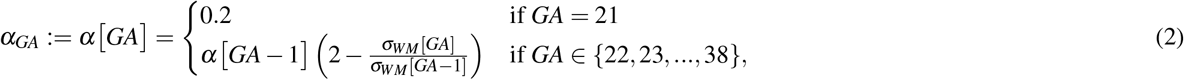

where *σ*_*WM*_ stands for the estimated standard deviation of WM intensity (see Figure 3-C). We empirically set *α*_21_ = 0.2, and iteratively compute *α* corresponding to the GA interval according to Equation 2 for the 17 remaining subjects of the STA. For continous GA estimation, *α*(·) is determined by shape-preserving piece-wise cubic interpolation for GA between 20.0 and 38.0 weeks. Given (2), we obtain the new T_1_ and T_2_ (see Figure 3-D) from the reference T_*k*_ as:

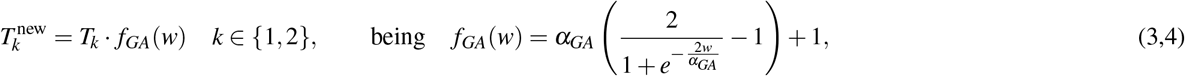

### Simulated datasets

High-resolution multi-tissue annotations from clinical experts are used as input models of the developing brain to simulate typical fetal brain FSE acquisitions, either at 1.5 T or 3 T^26^. For each subject, T2w series of 2D thick slices are simulated in the three orthogonal orientations (i.e., axial, coronal, sagittal) with respect to the fetal brain position. As routinely performed in clinical practice, two partially overlapping LR series are generated in every orientation for subsequent SR reconstruction of the fetal brain volume. Since the realism of interslice, 3D random rigid movements of the fetus encoded during k-space sampling achieved with FaBiAN v1.2^32^ has already been validated^26^, the dataset showcased in this work is generated without additional movement not to induce any bias in the qualitative evaluation of the realism of the generated LR series by the radiologists, but also to make it possible to align SR reconstructions from the simulated images to the corresponding ground truth label maps they originate from.

Table 1 reports the range of MR sequence parameters that determine the contrast, the geometry, and the resolution of representative clinical low-resolution series acquired in clinical routine on different MR systems (Half-Fourier Acquisition Single-shot Turbo spin Echo, HASTE for Siemens, Single-Shot FSE, SS-FSE for GE scanners), at either 1.5 or 3T, to screen the *in utero* developing brain. These parameters are therefore reproduced to simulate the T2w images of the fetal brain used throughout this study.

**Table 1.** Acquisition parameters at clinical magnetic field strength (1.5 and 3T) of two MR sequences (HASTE and SS-FSE) commonly used for fetal brain examination, and reproduced to simulate T2w images of the developing fetal brain.

| MR sequence | HASTE |  | SS-FSE |  |
| --- | --- | --- | --- | --- |
| Magnetic field strength (T) | 1.5 | 3 | 1.5 | 3 |
| Contrast |  |  |  |  |
| Effective echo time (ms) | 82-90 | 101 | 116-124 | 117-123 |
| Echo spacing (ms) | 4.08 | 10 | 10 | 10 |
| Echo train length | 134 | 126 | 224 | 224 |
| Excitation flip angle (°) | 90 | 90 | 90 | 90 |
| Refocusing pulse flip angle (°) | 149-180 | 180 | 150-180 | 150-180 |
| Geometry |  |  |  |  |
| Slice thickness (mm) | 3 | 3 | 3 | 3 |
| Slice gap (mm) | 0.3 | 0 | 0 | 0 |
| Phase oversampling (%) | 80 | 63 | 0 | 0 |
| Shift of the field-of-view (mm) | 0 | 0 | ±1.6 | ±1.6 |
| Resolution |  |  |  |  |
| Field-of-view in the readout direction (mm) | 360 | 350 | 240-300 | 240-320 |
| Base resolution (voxels) | 320 | 320 | 256 | 256 |
| Phase resolution (%) | 70 | 75 | 100 | 100 |
| Reconstruction matrix (voxels) | 320 | 640 | 512 | 512 |
| Zero-interpolation filling | none | yes | yes | yes |
| Acceleration technique |  |  |  |  |
| Acceleration factor | 2 | 2 | 1 | 1 |
| Reference lines | 42 | 24 | 0 | 0 |
| Noise |  |  |  |  |
| Mean | 0 | 0 | 0 | 0 |
| Standard deviation | 0.05 | 0.004 | 0.03-0.07 | 0.002-0.006 |

Besides non-standardized MR sequence parameters, only a few studies have investigated T_1_ and T_2_ properties of the developing brain, either *in utero*^38,39,41^ or in preterm newborns^37,40^. We determined mean reference T_1_^40,41^ and T_2_ values^37–39^ of GM and WM based on measurements performed at 1.5T, knowing that both T_1_ and T_2_ properties decrease across brain development (so from *in utero* to postnatal life) and that T_2_ > T_2_^*^. Since the biochemical composition of the CSF does not vary between childhood and adulthood, T_1_ and T_2_ values of this tissue were assumed to be similar in both populations, by extension also in fetuses. Besides, we extended these reference values to 3T by assuming that T_2_ relaxation time remains constant across clinical magnetic field strengths for a given structure, while T_1_ relaxation time increases by about 25% in GM and 10% in both WM and CSF^43–47^. As reported in Table 2, we then randomly generated T_1_ and T_2_ values in the range of mean T_*k*_ *±* standard deviation (SD) (*k* ∈ [[1, 2]]) to increase the diversity in tissue relaxation times across GA and within the three main classes modeled in the simulated images, as well as to account for measurement uncertainty in previous work.

**Table 2.** Range of fetal brain tissue properties simulated in this study at 1.5T and 3T.

| | $T_1$ (ms) | | $T_2$ (ms) |
| --- | --- | --- | --- |
|  | at 1.5T | at 3T | at 1.5T and 3T |
| Gray matter | $2200 \pm 150$ | $2750 \pm 150$ | $182 \pm 10$ |
| White matter | $2700 \pm 300$ | $2970 \pm 300$ | $285 \pm 15$ |
| Cerebrospinal fluid | 4000 | 4400 | 2000 |

## Data Records

The simulated LR series of the 78 subjects included in this study have been released on Zenodo (^48^) together with the corresponding brain masks and SR reconstructions. They are provided as compressed NIfTI images and organized in the Brain Imaging Data Structure (BIDS) format^49^. The Matlab (MathWorks, R2019a) and Python scripts used to generate this dataset have also been made publicly available (FaBiAN v2.0^34^). Moreover, the implementation of FaBiAN v1.2 has been slightly modified to be containerized into a docker image, and therefore facilitate the simulation of additional T2w MR images of the developing fetal brain from HR annotations. With the perspective of broadly disseminating this tool to the community, a similar docker image for FaBiAN v2.0 is available in DockerHub^50^.

### Technical Validation

#### Qualitative evaluation of the realism of the simulated low-resolution series

Figure 4 illustrates how locally adjusting T_1_ and T_2_ values within WM tissues enhances the resemblance of simulated FSE images with real clinical MR acquisitions of the fetal brain across maturation. It shows a comparison between synthetic LR series generated using both versions of FaBiAN from the segmented, SR-reconstructed fetal brain volumes of three representative subjects and the corresponding clinical images acquired at CHUV at 1.5T, in both healthy and pathological fetuses of 21, 31, and 33 weeks of GA respectively. The modulation of T_1_ and T_2_ properties according to the water/myelin content of WM tissues results in local variations of the MR contrast allows to better depict the complexity of WM and the underlying maturation processes. For instance, the migration of neurons from the germinal matrix to the cortex during the first two trimesters of gestation is reflected by a multilayer aspect of WM, which is finely captured in synthetic images generated using FaBiAN v2.0. In contrast, WM tissues appear highly homogeneous in the LR series simulated by FaBiAN v1.2 (see Figure 4-top row, in the coronal orientation).

**Figure 4.**
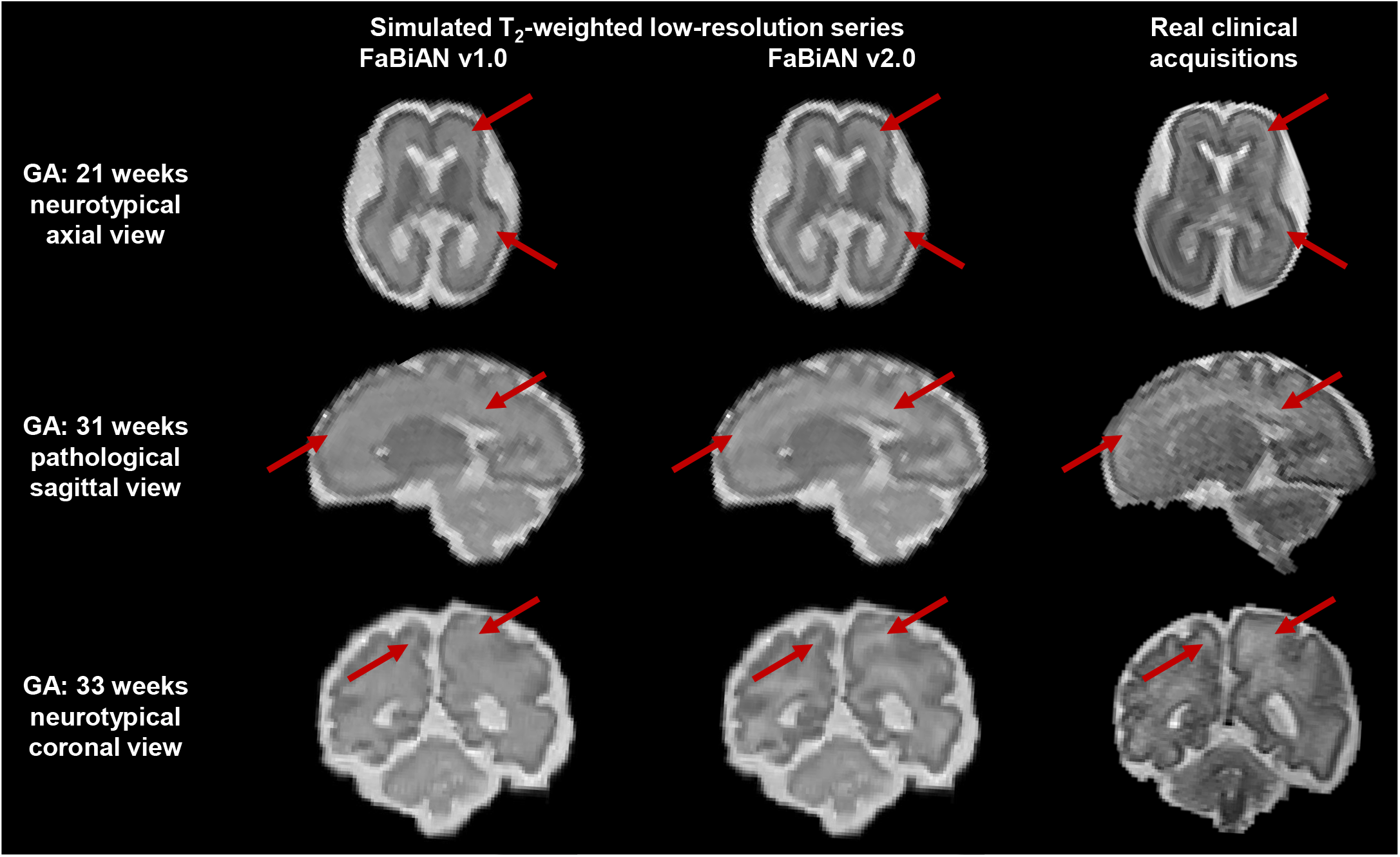
Comparison between synthetic low-resolution series generated in the different orthogonal orientations using FaBiAN v1.0 (left column) and v2.0 (middle column) and three representative clinical fast spin echo acquisitions (right column) at 1.5T in both healthy and pathological fetuses of 21 (top line), 31 (middle line), and 33 weeks (bottom line) of gestational age (GA) respectively scanned at CHUV. Red arrows highlight areas where white matter heterogeneities result in MR contrast variations that are representative of the ongoing maturation processes. These white matter changes are well captured by FaBiAN v2.0 simulations, which therefore look more realistic and closer to clinical acquisitions compared to images generated by FaBiAN v1.2.

##### Experimental design

A pediatric neuroradiologist and a neuroradiologist from CHUV with 17 and 14 years of experience respectively, provided independent, blind, qualitative assessment of the realism of the synthetic T2w LR series of the fetal brain generated for 29 subjects (16 neurotypical and 13 pathological) in the GA range of 20.1 to 34.8 weeks (27.0 *±* 4.20 weeks) by both the original version of the software (FaBiAN v1.2)^32^ and our new numerical WM model (FaBiAN v2.0)^34^. Young fetuses diagnosed with spina bifida prior surgery were excluded from this evaluation as the absence of extra-axial CSF substantially alters the quality of the clinical acquisitions and subsequent simulations. Simulations were run at 1.5T (25 subjects) and 3T (four subjects), the main difference being the higher spatial resolution and higher SNR that can be reached at 3T. High-quality LR series were simulated with high SNR and without motion not to bias the evaluation of the radiologists with corrupted images. Visual inspection and navigation throughout the different series of the fetal brain were made possible via ITK-SNAP^51^, displaying simulations in the same orientation plane but generated by both versions of the software in two different windows. Both experienced radiologists were asked to evaluate: i) which one of both series appears the most realistic (Experiment 1), ii) how realistic and close to MR images acquired in clinical routine every of these simulated series looks, taken independently, with a special focus on WM appearance, based on a five-point Likert scale (from 1: poorly realistic, to 5: highly realistic) (Experiment 2).

##### Experiment 1: Comparison between FaBiAN v1.2 and FaBiAN v2.0

This validation aims at comparing which version of our numerical phantom provides the most realistic LR series of the fetal brain throughout maturation in every orthogonal orientation. Figure 5 (top row) illustrates at which frequency every expert rated both versions of FaBiAN as providing the most realistic LR series compared to each other across development. Unanimously, FaBiAN v2.0 generated more realistic LR series over gestation than FaBiAN v1.2, for all subjects according to our neuroradiologist, respectively in more than 96.5% of the cases to our pediatric neuroradiologist. Interestingly, the three out of 87 series generated by FaBiAN v1.2 that they assessed as more realistic than the ones simulated by FaBiAN v2.0 corresponded to images of young fetuses (i.e., below 26 weeks of GA). At this early stage of development, the differences in the simulated images between both versions of FaBiAN may be more subtle to distinguish.

**Figure 5.**
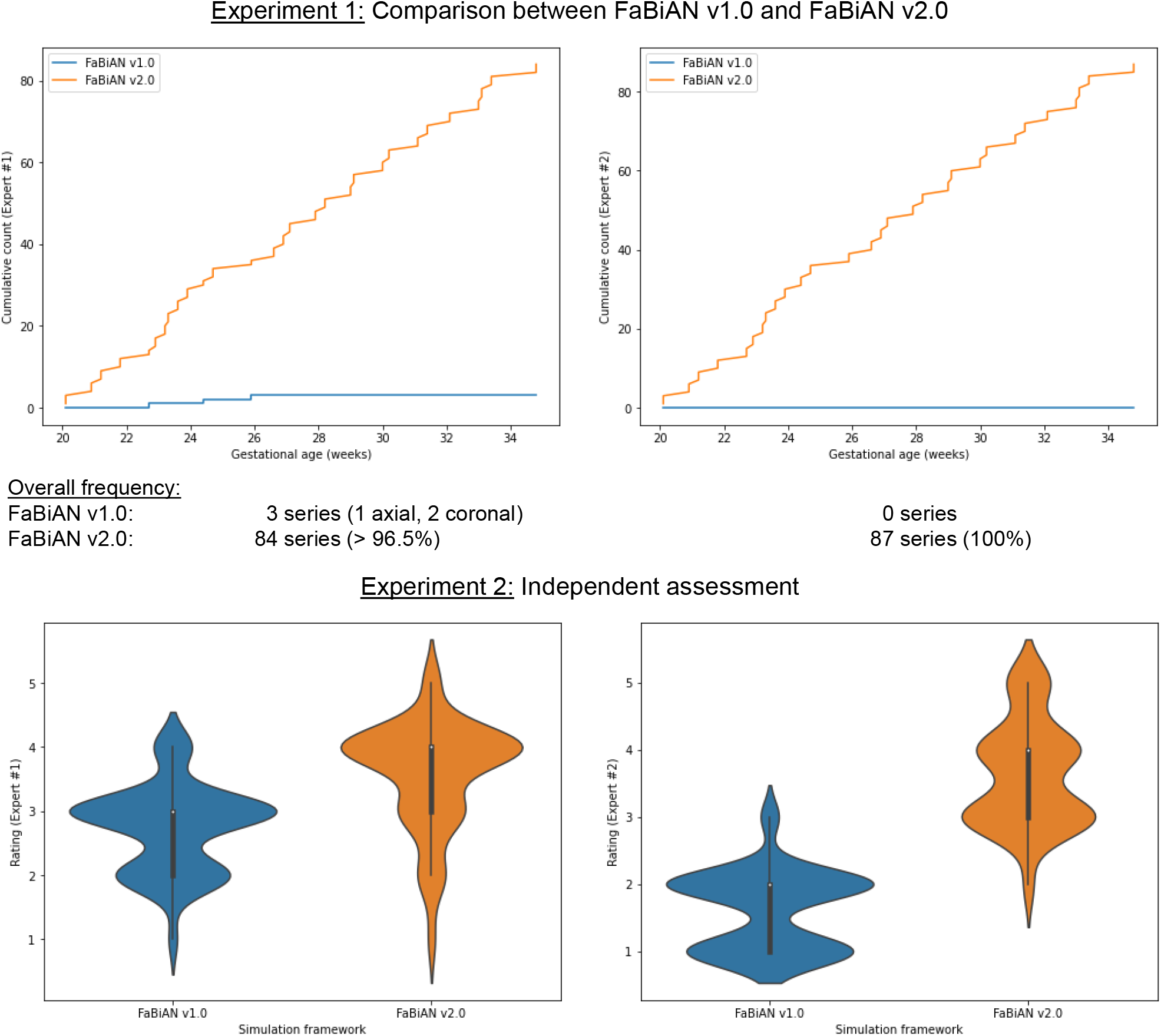
Qualitative evaluation of the realism of the low-resolution (LR) series simulated using FaBiAN v1.2 and FaBiAN v2.0 for 29 subjects, in the three orthogonal orientations, by two independent experts in pediatric neuroradiology (Expert #1) and in neuroradiology (Expert #2). Experiment 1 (top row) consisted in comparing images generated for every subject by both implementations of the simulation framework. The cumulative count of the most realistic synthetic LR series is represented over the gestational age, according to each expert, showing a net preference for FaBiAN v2.0 (between 96% and 100% of the images evaluated as the most realistic by Expert 1 and Expert 2 respectively) over FaBiAN v1.2 throughout fetal brain maturation. Experiment 2 (bottom row) further aimed at evaluating the realism of the simulated LR series independently. The violin plots show the distribution of the ratings (between 1-poorly realistic and 5-highly realistic) of every image by each expert. Overall, the LR series simulated by FaBiAN v2.0 were rated as more realistic than the ones generated using FaBiAN v1.2.

##### Experiment 2: Independent assessment

This experiment aims to independently evaluate the realism of the LR series of the developing fetal brain simulated by FaBiAN v1.2 and FaBiAN v2.0. The violin plots in Figure 5 (bottom row) display the distribution of ratings by each expert of all the synthetic LR series generated using both phantom versions. They show dispersity in the realism of the simulated images, although LR series generated by FaBiAN v2.0 were evaluated as the most realistic overall. These independent ratings of all the synthetic images provided, without comparison of both versions of the phantom to each other, further support the findings inferred from the first experiment, namely that our latest developments (FaBiAN v2.0) make it possible to generate even more realistic images than the original prototype (FaBiAN v1.2) by capturing local changes within WM tissues throughout maturation.

Both of these experiments validate that the proposed dataset generated using FaBiAN v2.0 provides highly realistic T2w MR images of the brain throughout *in utero* development, and not only in older fetuses where maturation processes are further advanced.

#### Quantitative comparison between simulated and original clinical images

This section aims at quantifying the similarity between simulated (FaBiAN v1.2, respectively FaBiAN v2.0) and real clinical data. Since SR reconstructions are of great value for further quantitative analysis of the developing fetal brain, and because manual annotations of these data are available, we compare the similarity between SR reconstructions from simulated and real cases (n=29 subjects), with a special focus on WM tissues.

Partially-overlapping orthogonal LR series (two in each acquisition plane) of the same 29 subjects inspected during qualitative evaluation by the radiologists were generated and combined to reconstruct an SR volume of the fetal brain of the corresponding clinical cases using two different pipelines, the Image Registration Toolkit SVRTK^18^ under Licence from Ixico Ltd., or MIALSRTK^19^), as in^16^. Overall, 21 subjects (nine neurotypical and 12 pathological in the GA range of 20.1 to 34.8 weeks (26.4 *±* 4.06 weeks) were reconstructed using SVRTK, eight subjects (seven neurotypical and one pathological in the GA range of 21.2 to 33.4 weeks (28.5 *±* 4.2 weeks) with MIALSRTK respectively. For every subject, mutual information (MI) was computed between the SR reconstructions from synthetic data (FaBiAN v1.2 against FaBiAN v2.0), and between the SR reconstructions from simulated (FaBiAN v1.2 or FaBiAN v2.0, respectively) and real clinical data, as a measure of similarity between them, with a focus on the WM mask. The higher the MI between two images, the closer they are.

Figure 6 quantifies to which extent SR reconstructions from simulated, respectively from real clinical acquisitions, are close to each other for the same subject and across GA. As shown by the left panel (blue dots), the differences between SR reconstructions from images generated by FaBiAN v1.2 and FaBiAN v2.0 are more pronounced in older fetuses, leading, for the same subject, to a decrease in MI between SR reconstructions from data simulated by FaBiAN v1.2 and FaBiAN v2.0, respectively, with GA. This trend can be explained by the growing proportion of WM in relation to other brain tissues over development, knowing that FaBiAN v2.0 accounts for local WM changes throughout brain maturation while WM appears rather uniform in simulations from the first prototype. The right panel of Figure 6 further supports these findings by highlighting an increase in MI between SR reconstructions from data simulated by FaBiAN v2.0 and the corresponding clinical subjects across development (green dots) whereas, on the contrary, MI between SR reconstructions from images generated by FaBiAN v1.2 and real subjects decreases with GA (red dots). Linear regressions on these data (see solid lines) corroborate that SR reconstructions of subjects older than 23 weeks of GA simulated by FaBiAN v2.0 show a greater similarity with real clinical data than their twins reconstructed from images generated by FaBiAN v1.2, especially as the fetus gets older. We, therefore, assume that local WM changes implemented by FaBiAN v2.0 throughout brain maturation enhance the realism of the simulated images by accounting for spatial and temporal heterogeneities within WM tissues.

**Figure 6.**
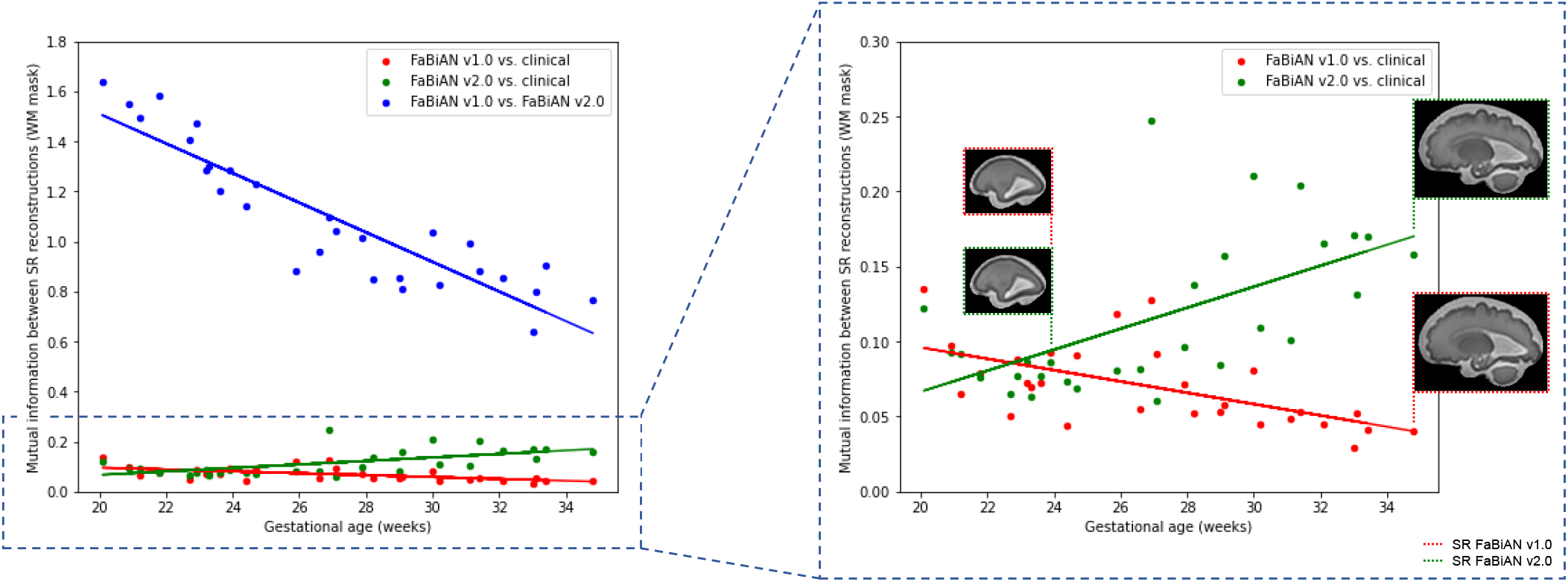
Mutual information (MI) between SR reconstructions from simulated data (FaBiAN v2.0 versus v1.2), and between SR reconstructions from simulated data (FaBiAN v1.2 or v2.0, respectively) and the corresponding SR reconstruction obtained from real clinical acquisitions across gestational age (GA) for the 29 subjects included in the qualitative evaluation by the radiologists, with a focus on the white matter mask. MI between SR reconstructions from simulated data (FaBiAN v2.0 versus v1.2) decreases with GA, showing a higher difference between the images generated by FaBiAN v2.0 and v1.2 at later GA, in other words when white matter changes linked to brain maturation become more obvious. The right panel of the Figure shows an enlargement of the plots that compare SR reconstructions from simulated and clinical data. SR reconstructions from data generated by FaBiAN v2.0 lead to an increase in MI across GA compared to images simulated using FaBiAN v1.2, thus demonstrating a higher similarity between the SR reconstructions from clinical subjects and data generated by FaBiAN v2.0 compared to images simulated using FaBiAN v1.2 as the fetus gets older.

#### Leveraging in silico data for automated fetal brain multi-tissue segmentation with deep learning

We expanded the technical validation of our proposed dataset as a valuable complement to scarce, annotated clinical data, in the context of fetal brain multi-tissue segmentation.

##### Methodology

###### Clinical data

Eighty-eight SR-reconstructed clinical subjects from the FeTA dataset^16^ were complemented with corresponding refined HR label maps^35,36^ that delineate eight classes: WM (excluding the corpus callosum), cortical GM, deep GM, extra-axial CSF, intra-axial CSF, brainstem, cerebellum, and corpus callosum. The dataset was split into training and test samples, including 70 (healthy: 27, pathological: 43) and 18 subjects (healthy: 7, pathological: 11), respectively, in the GA range of 20.0 to 34.8 weeks (27.1 *±* 3.62 weeks) and 20.9 to 33.1 weeks (26.9 *±* 3.53 weeks). Subjects were selected on a stratified random basis with disease condition and GA as factors.

###### In silico dataset

We generated realistic T2w MR images from the refined, clinical HR annotations based on the sequence parameters extracted from multiple LR series acquired in 15 representative subjects^26^, using our extended framework (FaBiAN v2.0) to account for WM heterogeneity and changes across maturation. For every subject, six partially overlapping 2D LR series were simulated in the three orthogonal orientations without motion and combined to reconstruct one single HR volume of the fetal brain using either SR reconstruction pipeline to match the corresponding clinical subject (either MIALSRTK v2.1.0, docker container^52^, or SVRTK^18^, using their default parameters).

###### Experimental design

Our study’s primary objective is to assess the practicality and efficacy of utilizing simulated data for training DL models in the context of segmentation tasks, prioritizing practical applications over the pursuit of a marginal, residual performance gain. To achieve this goal, we chose to employ the widely accessible nnU-Net implementation^53^, renowned for its robustness in similar tasks, as demonstrated by its strong performance in the FeTA Challenge 2021^17^. To evaluate the impact of simulated data, we conducted a comparative analysis, employing the same nnU-Net model while varying the input data. We initially trained a *baseline model* using 70 out of 88 subjects from the real dataset, including manual annotations (ground truth). Subsequently, we trained a series of models using varying proportions of SR images reconstructed from synthetic LR series, along with the corresponding ground truth annotations for each subject, intending to match the size and characteristics (e.g., individual anatomical features, tissue distribution, common acquisition parameters) of the training dataset. Specifically, we trained models with none (i.e., only SR reconstructions from synthetic data), six, nine, 12, 15, and 18 clinical SR images, combined with simulated data up to 70 subjects. In line with nnU-Net’s standard preprocessing, we performed skull stripping (SynthStrip, FreeSurfer^54^) on the images before inputting them to the different models.

###### Evaluation

The performance of fetal brain tissue segmentation models was assessed based on the Dice similarity coefficient (DSC^55,56^), which quantifies the overlap between the predicted segmentation and the ground truth manual tissue annotations. The performance metrics are reported for the test set (18 subjects) and are denoted as 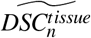, where *n* indicates the number of synthetic subjects used for training, and *tissue* consists of the following categories: WM, intra-axial CSF, cerebellum, extra-axial CSF, cortical GM, deep GM, brainstem, and corpus callosum.

##### Feasibility study

Our experiment evaluates whether automated segmentation of the developing fetal brain can be achieved with good performance without employing real data.

The boxplots in Figure 7-a) show that a model trained exclusively on synthetic images can achieve an acceptable overall performance in term of tissue segmentation 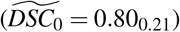, suggesting that simulated data contain sufficient semantics for the nnU-Net model to provide approximate annotations that could then be refined by a radiologist.

**Figure 7.**
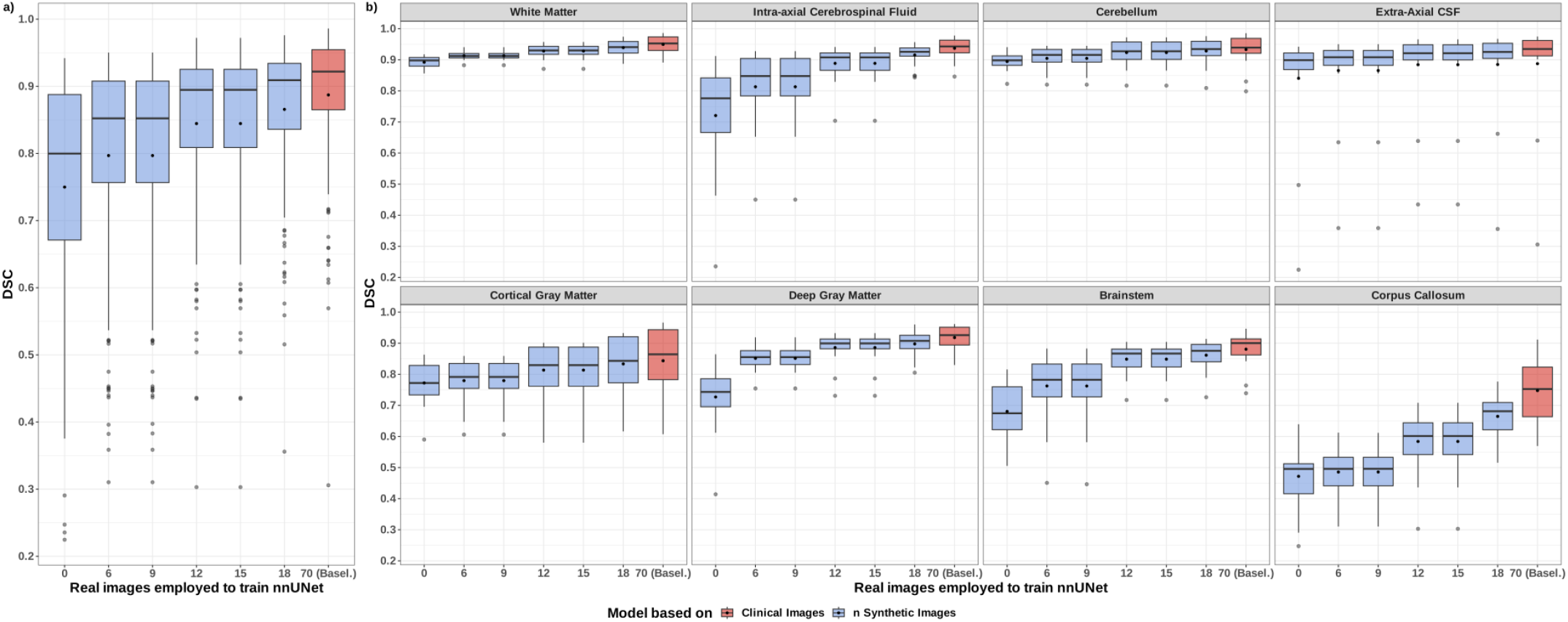
Dice similarity coefficient (DSC) scores for 18 test subjects during multi-tissue segmentation using nnU-Net with synthetic image training. Model performance is shown as the number of synthetic images decreases from 70 to zero: a) overall performance for all tissues and subjects, b) performance per tissue across all subjects.

Notably, the DSC substantially improves with the inclusion of a small number of real images in the training set (e.g., 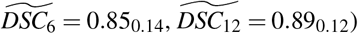, approaching the performance of a model trained solely on real images 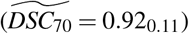. This observation highlights the potential for reducing the reliance on real data for training purposes, which is especially crucial in very specific or sensitive populations for which data are scarce.

Furthermore, it is important to note that the differences in model performance are not consistent across all tissues, as depicted in Figure 7-b). The most significant differences between models are primarily attributed to tissues that include complex structures, or to small regions of interest like the corpus callosum, which present challenges for accurate segmentation (with or without synthetic subjects). Conversely, all models exhibit similar performance when segmenting WM areas (excluding the corpus callosum;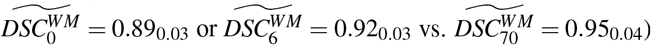, which have been a key focus of this work and the development of FaBiAN v2.0. This consistent reliability in the segmentation performance of WM further supports the potential for re-use of the dataset showcased in this work, as well as the validity of our new model of the developing fetal brain.

These findings provide valuable insights into the potential for using synthetic data to enhance the training of deep learning models for fetal brain tissue segmentation, particularly in scenarios where access to real data is limited or challenging.

## Code availability

The proposed framework is publicly is available on Zenodo (both dataset^48^ and software^34^). Synthetic images of the fetal brain throughout maturation were generated using MATLAB (MathWorks, R2019a). WM segmentation was performed using FMRIB’s Automated Segmentation Tool (FAST^42^, FMRIB Software Library, release 6.0.5.1). Simulated T2w MR images and corresponding annotations were reconstructed using the MIALSRTK pipeline^57^ (version 2.1.0, docker container).

## Acknowledgements

This work was supported by the Swiss National Science Foundation through grant 182602, and by the ProTechno Foundation. We acknowledge access to the facilities and expertise of the CIBM Center for Biomedical Imaging, a Swiss research center of excellence founded and supported by Lausanne University Hospital (CHUV), University of Lausanne (UNIL), Ecole Polytechnique Fédérale de Lausanne (EPFL), University of Geneva (UNIGE) and Geneva University Hospitals (HUG). UPC 389 activity in this project has been supported by PID2020 116907RBI00, funded by MCIN AEI 10.13039 501100011033.

## Author contributions statement

H.L., P.M.G., O.E., and M.B.C. conceived the experiments. A.B.F. and H.L. modified the framework based on their expertise and the feedback from O.E., F.M., and M.B.C.. A.B.F. prepared the preliminary work on which this manuscript relies under the supervision of H.L. and M.B.C.. H.L. simulated all images used throughout this work. M.K. and V.D. assessed the realism of the simulated images. P.M.G. trained and tested the 3D nnU-Net for automated fetal brain multi-tissue segmentation. H.L., A.B.F., and P.M.G. processed the results. H.L., P.M.G., O.E., and M.B.C. analyzed the results. All authors reviewed the manuscript.

## Competing interests

The authors declare no competing interests.

## Notes

https://doi.org/10.5281/zenodo.10940427

https://hub.docker.com/r/petermcgor/fabian-docker

https://github.com/Medical-Image-Analysis-Laboratory/FaBiAN

### Competing Interest Statement

The authors have declared no competing interest.

